# Poly(ADP-ribose) Polymerase 1 Deficiency Attenuates Amyloid Pathology, Neurodegeneration, and Cognitive Decline in a Familial Alzheimer’s Disease Model

**DOI:** 10.1101/2025.09.06.674313

**Authors:** Aanishaa Jhaldiyal, Manisha Kumari, Trupti Tripathi, Mohammed Repon Khan, Justin Wang, Lauren Guttman, Devanik Biswas, Abhishek Pasupuleti, Akansha Aggarwal, Shraddha Pandya, Shih-Ching Chou, Nikhil Panicker, Abhay Monghekar, Marilyn Albert, Lynn M. Bekris, James B. Leverenz, Tae-In Kam, Ted M. Dawson, Valina L. Dawson

**Affiliations:** Neuroregeneration and Stem Cell Programs, Institute for Cell Engineering, The Johns Hopkins University School of Medicine, Baltimore, MD, 21205, USA; Department of Neurology, The Johns Hopkins University School of Medicine, Baltimore, MD, 21205, USA; Department of Physiology, Pharmacology and Therapeutics, The Johns Hopkins University School of Medicine, Baltimore, Maryland, 21205 USA; Department of Biochemistry and Molecular Biology, Bloomberg School of Public Health, Johns Hopkins University School of Medicine, Baltimore, Maryland 21205, USA; Department of Laboratory Medicine and Pathology, University of Washington, Harborview Research and Training Building, 300 Ninth Ave, Box 359665, Seattle, WA 98104; Lerner Research Institute, Genomic Medicine, Cleveland Clinic, Cleveland, OH 44195, USA; Lou Ruvo Center for Brain Health, Neurological Institute, and Department of Neurology, Cleveland Clinic, Cleveland, OH 44195, USA; Geriatric Research Education and Clinical Center, VA Puget Sound Health Care System (S-182), 1660 South Columbian Way, Seattle, WA 98108, USA; Department of Neurology, University of Washington, 1959 NE Pacific St, Seattle, WA 98195, USA; Solomon H. Snyder Department of Neuroscience, The Johns Hopkins University School of Medicine, Baltimore, MD, 21205, USA; Cleveland Clinic Research, Cleveland Clinic, Cleveland, OH 44195, USA; Department of Brain and Cognitive Sciences, Korea Advanced Institute of Science and Technology (KAIST), Daejeon 34141, Republic of Korea; Graduate School of Stem Cell and Regenerative Biology, Korea Advanced Institute of Science and Technology (KAIST), Daejeon 34141, Republic of Korea; Passed away prior to completion of the manuscript

**Keywords:** Poly(ADP-ribose) (PAR), PARP1, Alzheimer’s disease, Amyloid beta (Aβ), Neurodegeneration

## Abstract

Poly(ADP-ribose) (PAR) polymerase-1 (PARP1) has been implicated in DNA damage responses and neuroinflammation in Alzheimer’s disease (AD), yet its role in amyloid-β (Aβ) pathology remains unclear. Here, we show that PARP1 activation drives Aβ pathology and neurodegeneration. Using a sensitive ELISA, we observed significantly elevated PAR levels in the cerebrospinal fluid (CSF) of patients with mild cognitive impairment (MCI) and AD compared to controls. *In vitro*, oligomeric Aβ_1-42_ activated PARP1 and induced DNA damage, while genetic or pharmacological inhibition of PARP1 conferred neuroprotection. *In vivo*, PARP1 knockout in the 5XFAD mouse model of amyloidosis led to reduced amyloid plaque burden, preserved synaptic and neuronal integrity, attenuated glial activation and neuroinflammation, and rescued cognitive deficits. Mechanistically, PARP1 deficiency decreased amyloid precursor protein (APP) and BACE1 levels, altered γ-secretase complex composition, and enhanced Aβ degradation via neprilysin. These findings position PARP1 as a critical mediator of Aβ toxicity and neurodegeneration, suggesting its inhibition as a promising therapeutic strategy for AD.

**Significance Statement:** Our study identifies poly(ADP-ribose) (PAR) as an elevated biomarker in the cerebrospinal fluid of patients with mild cognitive impairment and Alzheimer’s disease, correlating with established markers of amyloid pathology. We demonstrate that PARP1, the enzyme responsible for PAR synthesis, is activated by neurotoxic Aβ_1-42_ and mediates neuronal death, amyloid plaque formation, neuroinflammation, and cognitive deficits in a mouse model of AD. Importantly, genetic ablation of PARP1 not only protects neurons from Aβ toxicity but also reduces amyloid burden by suppressing Aβ production and enhancing its degradation. These findings highlight PARP1 as a critical regulator of amyloid pathology and neurodegeneration, and suggest that PARP1 inhibition may offer a promising therapeutic avenue for Alzheimer’s disease by simultaneously targeting multiple pathogenic mechanisms.

## Introduction

Poly (ADP-ribose) polymerase 1 (PARP1) activation leads to neuronal cell death following glutamate excitotoxicity and stroke (1–4) and neurodegeneration in Parkinson’s disease (PD) (5–7). This form of cell death is designated parthanatos, and is distinct from other types of cell death such as apoptosis and necrosis (8, 9). Interference of each step in the parthanatic cascade has conferred neuroprotection in a variety of models (10–12) . It has generally been thought that PARP1 activity and parthanatos primarily play a role in acute cellular injury such as stroke and/or glutamate excitotoxicity. The view was changed with our recent report that PARP1 activity and parthanatos contribute to neurodegeneration in a chronic animal model of PD in which we discovered that degeneration of dopamine (DA) neurons in the substantia nigra pars compacta (SNpc) in the pathologic α-synuclein model of PD generated by intrastriatal injections of α-synuclein preformed fibrils (α-syn PFF) was primarily driven by PARP1 activity and parthanatos (5, 13, 14). Activation of macrophage migration inhibitory factor (MIF) nuclease activity, the final executioner of parthanatos drives pathology in three orthogonal models of PD further implicating parthanatos in a chronic neurodegenerative disorder (14). In postmortem human AD brain there is evidence of elevated PARP1 activity where Love et al., reported that product of PARP1 activity, Poly (ADP-ribose) (PAR) was elevated in the frontal and temporal cortices of AD patients versus controls (15). PARP1 polymorphisms may also decrease the risk and severity of AD (16).

However, the precise role of parthanatos in driving neuronal loss in AD is unknown. There are several transgenic models of AD (17). The 5XFAD transgenic mouse model has been extensively used to study amyloid β (Aβ) pathology in mice. The 5XFAD transgenic mice co-express a total of five FAD mutations [APP K670N/M671L (Swedish) + I716V (Florida) + V717I (London) and PS1 M146L+ L286V] (18). In addition to exhibiting memory impairment the 5XFAD transgenic mouse is one of the few Aβ animal models that exhibits significant neuron loss (19). By 9 months of age 5XFAD mice exhibit visible loss of large pyramidal neurons in cortical Layer 5 and the subiculum. Thus, 5XFAD mice allow for the investigation of whether parthanatos contributes to driving neuronal loss in the context of pathologic Aβ.

We crossed the 5XFAD mice with PARP1^-/-^ mice and report that the absence of PARP1 significantly reduces the neuronal loss, Aβ plaque load and cognitive deficits in the 5XFAD mice. This decrease in pathology was also accompanied by changes in markers of neuroinflammation and attenuation of glial pathology. Moreover, we reveal PARP1’s involvement in altering amyloid precursor protein (APP) metabolism, suggesting the enzyme plays a broader role in the pathogenesis of AD beyond its influence on neuronal loss.

## Results

### Increased levels of PAR in the cerebrospinal Fluid (CSF) of patients with mild cognitive impairment (MCI) and AD

A sensitive enzyme-linked immunosorbent assay (ELISA) for PAR (5) was used to determine whether PAR is elevated in the cerebrospinal fluid (CSF) of patients with MCI or AD compared to controls. PAR levels were significantly elevated in patients with MCI and AD versus controls in two independent patient cohorts (Fig. 1 *A, D,* Table S1). The Aβ42/40 ratio progressively decreases along the Alzheimer’s continuum from MCI to dementia (20). Accordingly, we observed the same trend in our CSF samples (Fig. 1 *B, E*). We also performed Pearson correlation analysis to examine the relationship between Aβ42/40 and PAR. In both cohorts, we found a negative correlation between PAR levels and the Aβ42/40 ratio in MCI samples (Fig. 1 *C, F*). Collectively, these data revealed increased CSF PAR in both MCI and AD, suggesting a potential role for PAR and PARP1 in AD pathogenesis.

**Figure 1.**
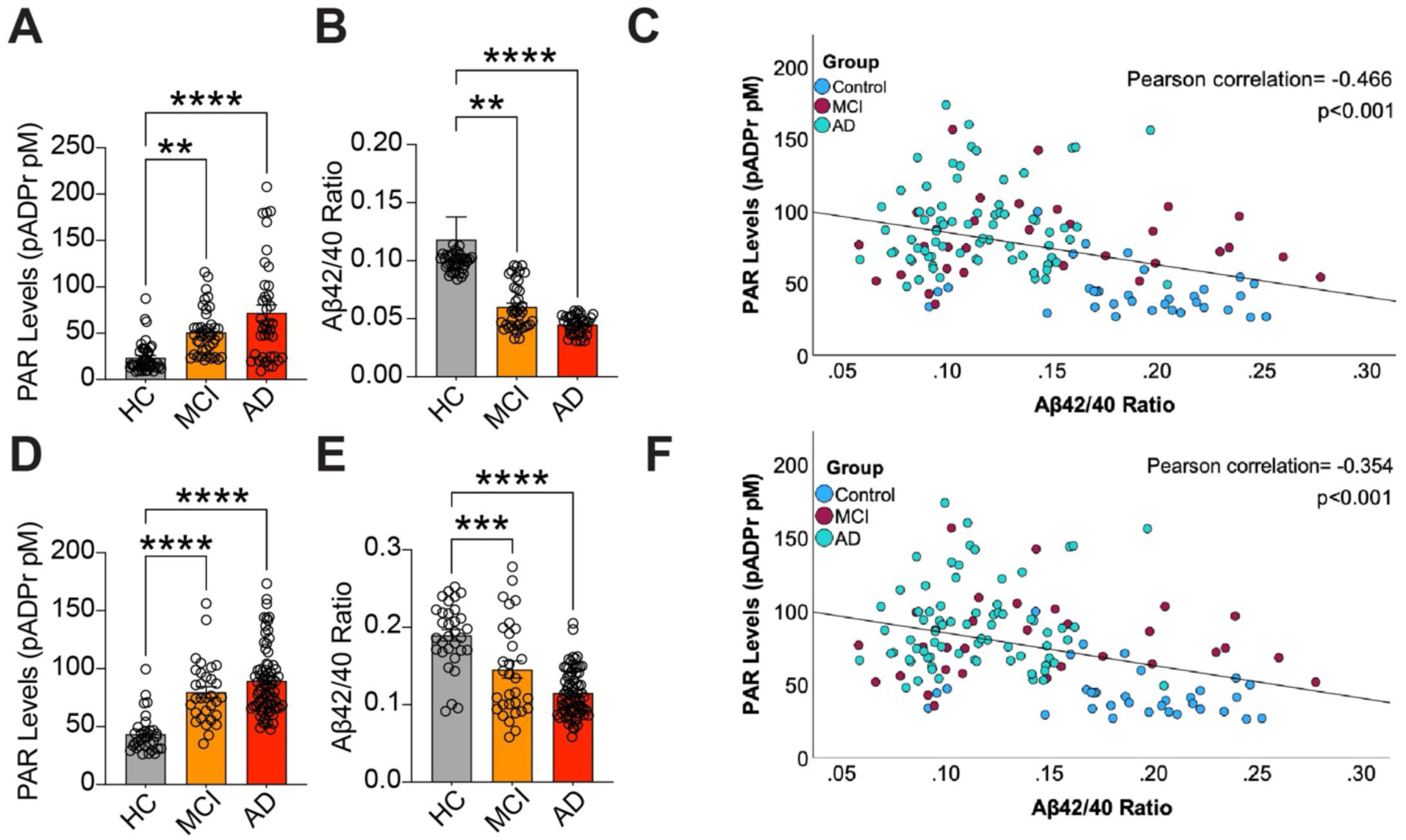
Poly (ADP-ribose) (PAR) levels are increased in patients with mild cognitive deficit (MCI) and Alzheimer’s Dementia (AD). A) PAR levels from CSF samples from Johns Hopkins Medicine of patients with MCI (n = 40) and AD (n=40) versus healthy control (HC) (n = 40) as determined by PAR ELISA. Bars represent means ± SEM. One-way analysis of variance (ANOVA) was followed by Tukey’s post hoc test. B) Aβ42/40 ratio with advancing stages of AD (HC>MCI>AD) of CSF samples from Johns Hopkins Medicine. Bars represent means ± SEM. One-way analysis of variance (ANOVA) was followed by Tukey’s post hoc test. C) Scatter plot of the correlation between CSF PAR (from Johns Hopkins Medicine) with Aβ40/42 ratio in HC, MCI and AD. Correlation coefficients (r) and p-values are from Pearson’s correlation analysis. D) PAR levels in CSF of healthy controls (HC) (n=33), patients with MCI (n=31) and patients with AD (n=74) as determined by PAR ELISA. Bars represent means ± SEM. One-way analysis of variance (ANOVA) was followed by Tukey’s post hoc test. E) Aβ42/40 ratio of CSF samples from Cleveland Clinic in advancing stages of AD (HC>MCI>AD). Bars represent means ± SEM. One-way analysis of variance (ANOVA) was followed by Tukey’s post hoc test. F) Scatter plot of the correlation between CSF PAR (from Cleveland Clinic) with Aβ40/42 ratio in HC, MCI and AD. Correlation coefficients (r) and p-values are from Pearson’s correlation analysis.

### Oligomeric Aβ_1-42_ activates PARP1, and genetic deletion and pharmacological inhibition of PARP1 protects against Aβ_1-42_ neurotoxicity

To explore a link between AD pathogenesis and PARP1, oligomeric Aβ species (Fig. S1*A*) were added to primary cortical neuron cultures to determine if PARP1 becomes activated. Either Aβ_1-40_ (1 μM) or Aβ_1-42_ (1 μM) was applied to primary cortical neurons and PARP1 activity was assessed 24 h later via PAR immunoblot analysis (Fig. 2*A, B*). Aβ_1-42_ administration led to a significant increase in PAR immunoreactivity while Aβ_1-40_ had no effect (Fig. 2*A, B*). We also evaluated the presence of γ-H2AX, a marker of DNA damage, which is concurrently elevated with PARP1 activity (Fig. 2*A,C*) (21).

**Figure 2.**
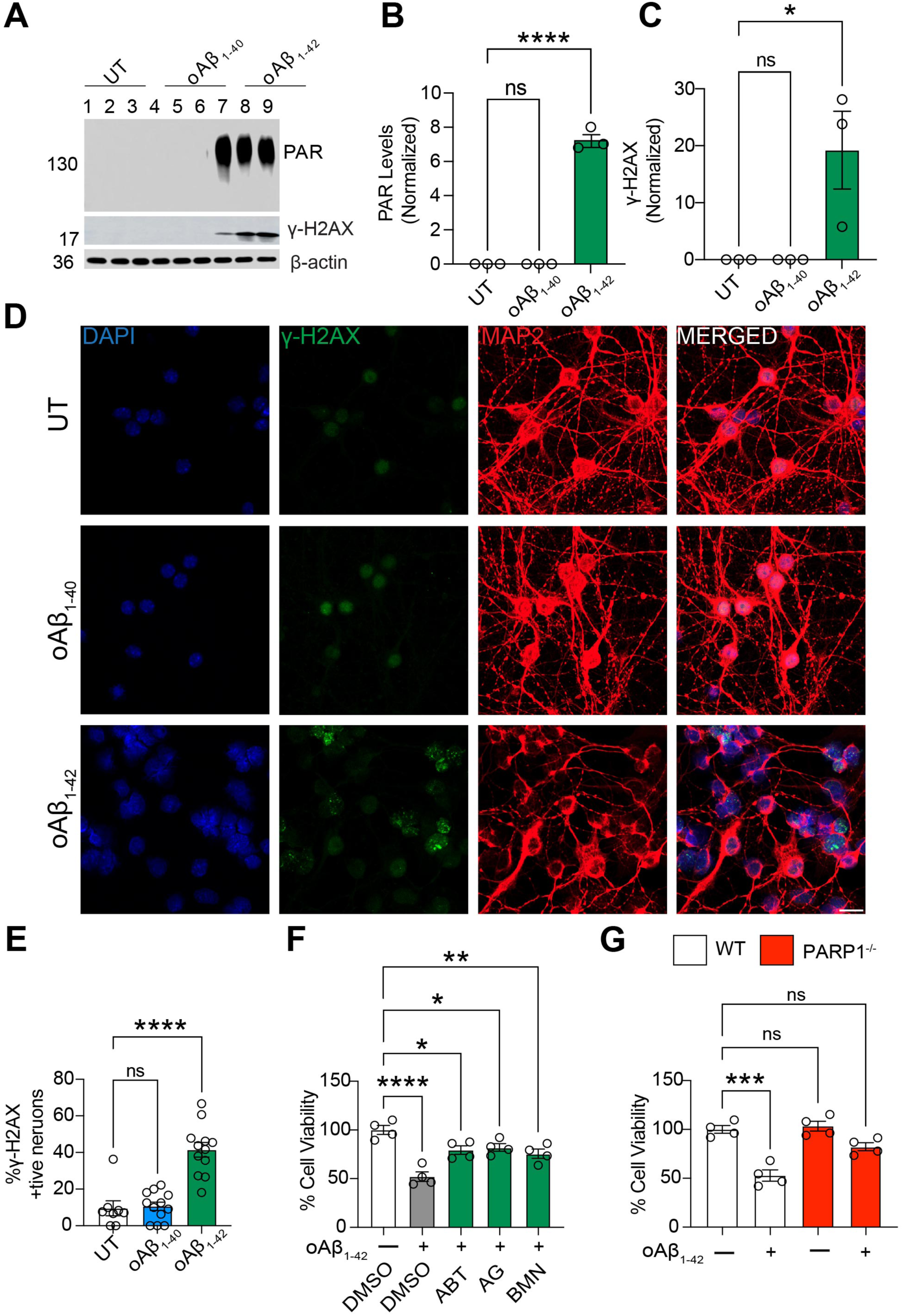
Oligomeric Aβ_1-42_ (oAβ_1-42_) treatment causes PAR activation in primary cortical neurons. A, B, C) Representative (A) immunoblot and quantification of (B) PAR and (C) γ-H2AX levels in primary cortical neurons untreated (UT) or treated with 1 µM oAβ_1-40_ or oAβ_1-42_ for 24 h. Bars are means ± SEM (n=3). One-way analysis of variance (ANOVA) followed by Tukey’s post hoc test. D, E) γ-H2AX–positive nuclei treated with 1 µM oAβ_1-42_ or 1 µM oAβ_1-40_ or in untreated (UT) controls. (D) Representative γ-H2AX immunofluorescence. Scale bar 20 µm.(F) Quantification of γ-H2AX–positive nuclei in UT, 1 µM oAβ_1-40,_ and 1 µM oAβ_1-42_ cultures. Bars represent mean ± SEM (n = 12). One-way ANOVA with Tukey’s post hoc test. G) Alamar blue cell viability assay from primary cortical neurons pre-incubated with 1 µM of ABT-888, AG-014699, or BMN 673 for 1 h and further incubated with 1 µM oAβ_1-42_ for 2 days. Bars represent mean ± SEM. Two-way ANOVA followed by Tukey’s post hoc test (n=4). H) Alamar blue cell viability assay from WT or PARP1 KO primary cortical neurons and further incubated with 1 µM oAβ_1-42_ for 2 days. Bars represent mean ± SEM. Two-way ANOVA followed by Tukey’s post hoc test (n=4).

Accompanying the increase in PAR levels was a statistically significant increase in γ-H2AX (Fig. 2A, *C*). γ-H2AX immunoreactivity was also significantly increased in the nuclei of cortical neurons treated with Aβ_1-42_ for 24 h (Fig. 2D*, E*). Oligomeric Aβ_1-42_ (1 μM) administration significantly increased PAR levels as early as 8 h, and there were further significant increases at 24 h and 48 h (Fig. S1*B, C*). A significant and steady increase in γ-H2AX was also observed along these time points (Fig S1*B, D*). These results indicate that oligomeric Aβ_1-42_ damages DNA and is an activator of PARP1.

The role of PARP1 in oligomeric Aβ_1-42_ toxicity was assessed by the Alamar blue cell viability assay (Fig. 2*F*). Oligomeric Aβ_1-42_ toxicity was significantly reduced in wild type (WT) neurons in the presence of PARP inhibitors ABT-888 (veliparib), AG-014699 (rucaparib), or BMN 673 (talazoparib) at 1 μM (Fig. 2*F*). ABT-888 (1 μM) also significantly protected WT neurons from cell death by oligomeric Aβ_1-42_ (Fig. S1 *E, F*), assessed using Propidium Iodide (PI) staining.. PARP1^-/-^ neurons were also significantly resistant to oligomeric Aβ_1-42_ neurotoxicity compared to WT neurons (Fig. 2*G*).

Additionally, oligomeric Aβ_1-42_ failed to produce PAR in PARP1^-/-^ neurons (Fig. S1G). Together, these findings demonstrate that PARP1 activation contributes to oligomeric Aβ_1-42_ –induced neurotoxicity, and that both pharmacological inhibition and genetic ablation of PARP1 confer significant neuroprotection.

### The absence of PARP1 reduces the Aβ plaque load and prevents neuronal cell death in 5XFAD mice

The role of PARP1 in Aβ pathology, neurobehavior, and neurodegeneration was assessed using 5XFAD transgenic mice (18). 5XFAD mice were crossed to PARP1^-/-^ mice to generate 5XFAD/PARP1^+/-^ mice, which were then crossed with PARP1^+/-^ mice. Approximately 20 male and female mice of WT, 5XFAD, PARP1^-/-^ and 5XFAD/PARP1^-/-^ were generated and aged to nine months (Fig. S2*A*). Agarose gel and immunoblot analysis confirmed the absence of PARP1 in 5XFAD/ PARP1^-/-^ and PARP1^-/-^ mice (Fig. S2*B, C*). APP mRNA level as determined by RT-PCR was significantly upregulated in 5XFAD mice compared to WT and PARP1^-/-^ mice. There was no significant difference in the APP mRNA level in 5XFAD versus 5XFAD/ PARP1^-/-^ mice (Fig. S2*D*).

5XFAD/PARP1^-/-^ animals exhibited a ∼50% reduction in Thioflavin S–positive plaque area compared with age-match littermate 5XFAD controls (Fig. 3A,B). To assess whether this decrease in plaque burden corresponded to changes in Aβ species, human Aβ_1-40_ and Aβ_1-42_ levels were measure by ELISA. In 5XFAD/PARP1^-/-^ mice, levels of Aβ_1-40_ and Aβ_1-42_ were significantly lower in TBS-soluble, TBS + Triton X-100–soluble, and 70% formic acid–soluble fractions relative to 5XFAD mice (Fig. 3 C,D). To confirm the reduction in aggregated amyloid species, we performed immunoblot analysis of high– molecular–weight Aβ aggregates isolated by sucrose cushion ultracentrifugation. This revealed elevated levels of aggregated Aβ and PAR in 5XFAD mice compared to 5XFAD/PARP1^⁻/⁻^ and littermate controls (WT and PARP1^⁻/⁻^) mice (Fig. 3*E-G*).

**Figure 3.**
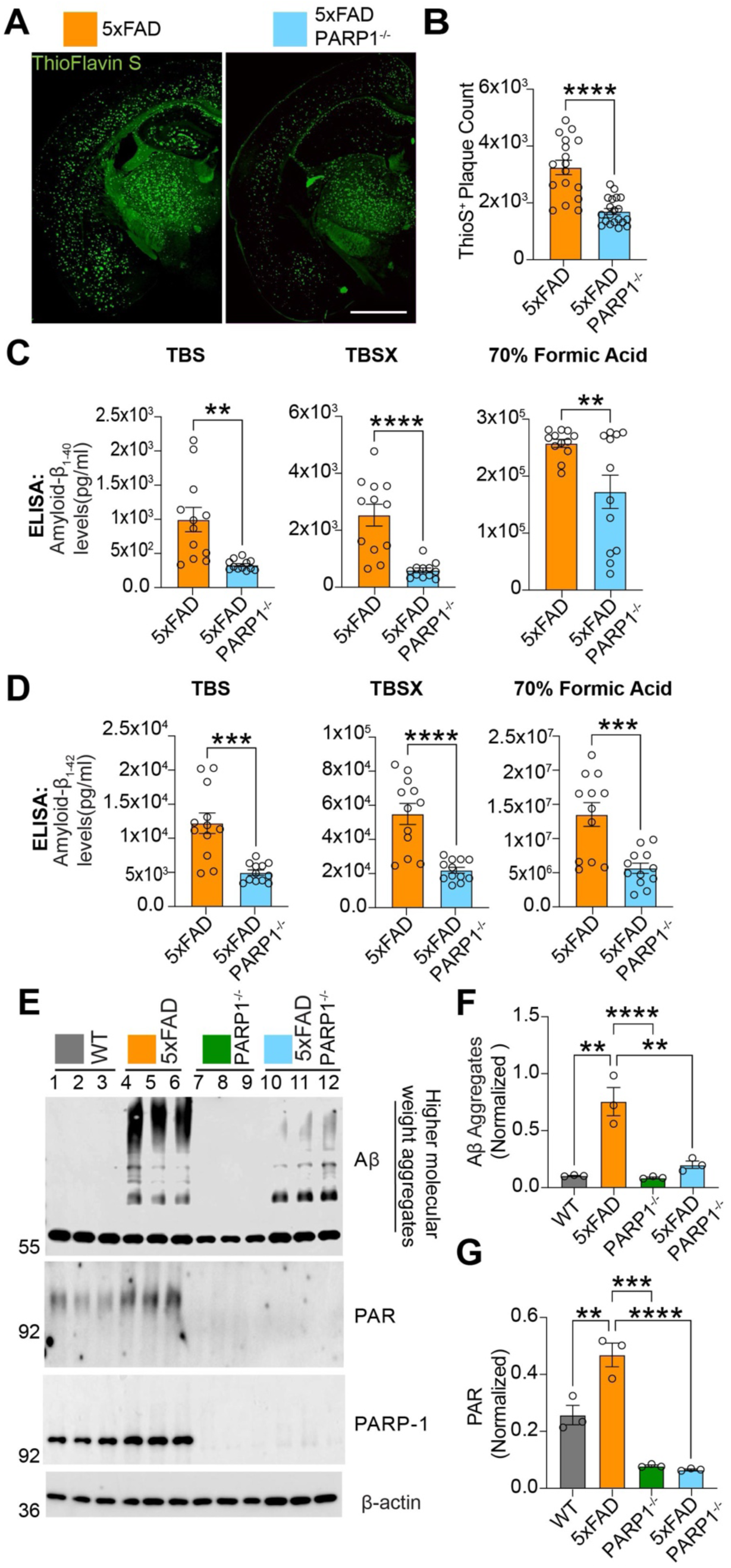
PARP1 deletion decreases Aβ plaque neuropathology in 5XFAD mice. (A) Immunofluorescence images and (B) quantification of Thioflavin S staining of amyloid-β plaques in 5XTg FAD and 5XFAD/PARP1^-/-^. Scale Bar 1000 µm. Bars are means ± SEM. Unpaired Student’s t test (n = 10). ELISA quantification of (C) Aβ_1-40_ and (D) Aβ_1-42_ levels in 5XFAD and 5XFAD/PARP1^-/-^. Bars are means ± SEM. Unpaired Student’s t test (n = 12). (E) Immunoblot and (F) quantification of extracted aggregate Aβ and (G) PAR from WT, PARP1^-/-^ , 5XFAD and 5XFAD/PARP1^-/-^mice. Immunoblot shows Aβ aggregates and PAR levels. Bars are means ± SEM. Two-way ANOVA followed by Tukey’s post hoc test (n =3).

Accompanying the reduction in Aβ core plaques was a preservation of synaptic density as assessed by PSD95 immunoreactivity density near Aβ core plaques labeled by 4G8 immunoreactivity (Fig. 4*A, B*). The accumulation of Aβ into fibrillar amyloid plaques causes damage to nearby neuronal structures, leading to the development of enlarged, swollen axons and dendrites around the plaques, a phenomenon known as neuritic dystrophy. These swollen neurites contain aggregates of several proteins, such as RTN-3, which can serve as markers for this dystrophic process (22, 23). RTN-3 levels were reduced in amyloid plaque–rich regions of 5XFAD/PARP1^-/-^ mice compared with 5XFAD controls. (Fig. 4*C,D*).

**Figure 4.**
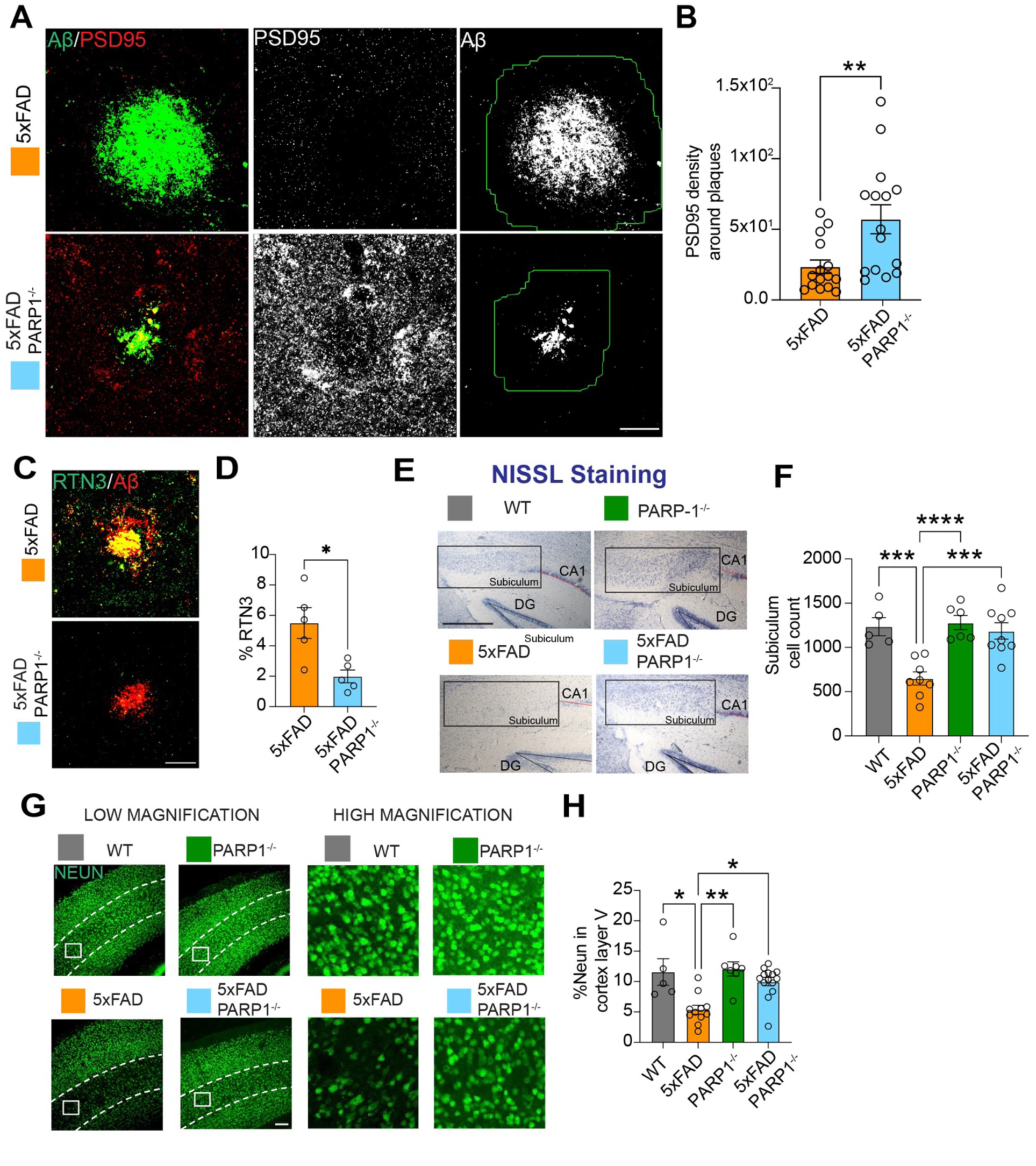
PARP1 deletion increases neuronal survival and synaptic activity in 5XFAD mice. A, B) Immunofluorescence images and quantification of PSD95 density surrounding amyloid-β plaques in 5XFAD and 5XFAD/PARP1⁻/⁻ mice. Cumulative PSD95 density was measured within a 30 µm radius of each plaque. Scale Bar 40 µm. Unpaired Student’s t test (n = 15). C, D) Immunofluorescent images and quantification of RTN3 levels around amyloid-β plaques in 5XFAD and 5XFAD/PARP1⁻/⁻ mice. Scale Bar 40 µm. Unpaired Student’s t test (n = 5). E, F) Nissl images and quantification of WT, PARP1^-/-^, 5XFAD and 5XFAD/PARP1⁻/⁻ mice.. Scale Bar 400 µm. Bars are means ± SEM. Two-way ANOVA followed by Tukey’s post hoc test (n = 4-10). G, H) NeuN staining immunofluorescence images and quantification of cortex layer V in WT, PARP1^-/-^, 5XFAD and 5XFAD/PARP1^⁻/⁻^ mice. Scale Bar 40µm. Bars are means ± SEM. Two-way ANOVA followed by Tukey’s post hoc test (n = 5-7).

In 5XFAD mice the subiculum and cortical layer V are areas with the most severe amyloidosis and neuron loss that begins at about 6 months of age (18, 19). Non-biased stereological cell counting revealed that neuronal cell loss was prevented in the 5XFAD/PARP1^-/-^ mice as assessed by Nissl stain in the subiculum of the hippocampus (Fig. 4*E, F*). NeuN immunostaining intensity was significantly preserved in layer V of the cerebral cortex in 5XFAD/PARP1^-/-^ mice compared to 5XFAD mice (Fig. 4*G, H*).

### Attenuation of markers of glial pathology and neuroinflammation in PARP1^-/-^ mice

Hippocampal brain sections of WT, 5XFAD, PARP1^-/-^ and 5XFAD/PARP1^-/-^ were immunostained with the microglia marker, ionized calcium-binding adaptor molecule 1 (IBA1), and the astrocyte marker, glial fibrillary acidic protein (GFAP). 5XFAD/PARP1^-/-^ mice exhibited a significant downregulation of astrogliosis (Fig. 5A, B) and microgliosis (Fig. 5A, C). The mRNA for the neuroinflammatory markers, TNF-α, C1q, IL-1β, IL-6 and C3 mRNA, were significantly upregulated in 5XFAD mice, while they were significantly decreased in 5XFAD/PARP1^-/-^ mice (Fig. 5D-H). Thus, our data suggest that loss of PARP1 reduces microglia and astrocyte activation.

**Figure 5.**
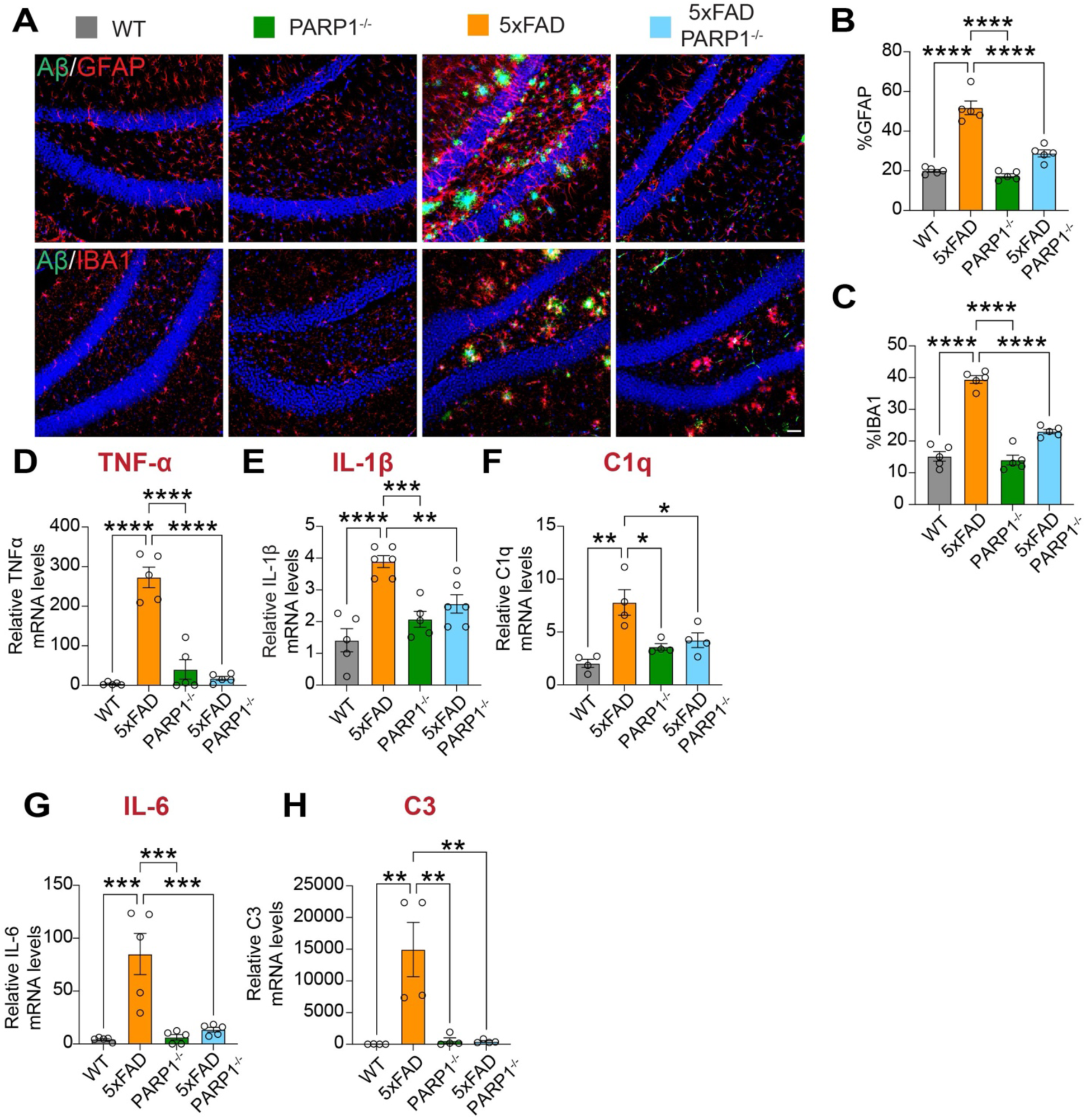
5XFAD/PARP1^-/-^ mice exhibit reduced gliosis. A-D) Immunofluorescent images and quantification of IBA1 and GFAP levels in WT, PARP1^-/-^, 5XFAD and 5XFAD/PARP1^-/-^ . Scale Bar 40 µm. Bars are means ± SEM. Two-way ANOVA followed by Tukey’s post hoc test (n = 5). D-I) mRNA levels of TNF-α, IL-1β, C1q, IL-6 and C3, as determined by RT-PCR. Bars represent mean ± SEM. Two-way ANOVA followed by Tukey’s post hoc test (n=5).

### Neurobehavioral deficits are reduced in 5XFAD mice lacking PARP1

Neurobehavior of WT, 5XFAD, PARP1^-/-^ and 5XFAD/PARP1^-/-^ was assessed via the Morris water maze, Y-Maze, and open field tests (Fig. 6). Spatial learning and memory were assessed by the Morris water maze task at 9 months. 5XFAD mice exhibited a significant decline in performance compared to WT and PARP1 KO mice. In contrast, 5XFAD/PARP1^-/-^ showed improved spatial and long-term memory across the 5-_day training period (__Fig. 6*A*_*_)._* Probe trials conducted 24 hours after the last training trial demonstrated that the 5XFAD/PARP^-/-^ mice spent significantly more time in the target quadrant than the 5XFAD mice (Fig. 6C), traveled a greater distance in the target quadrant (Fig. S3A), and had more entries in the target quadrant (Fig. S3B). _The Y-maze_ test was used to assess short-term or working memory. The percentage of alternating behaviors in the 5XFAD mice was significantly lower than that of WT, and PARP1^-/-^ mice (Fig. 6*D-F*). The 5XFAD/PARP1^-/-^ mice significantly prevented the deficits observed in 5XFAD mice. The open field test was performed to evaluate locomotor and anxiety-like behaviors. 5XFAD mice traveled significantly greater distances and spent significantly more time in the periphery than the WT, and PARP1^-/-^ mice (Fig. S3*C-E).* 5XFAD/PARP1^-/-^ did not exhibit anxiety like behavior (Fig. S3*C-E)*.

**Figure 6.**
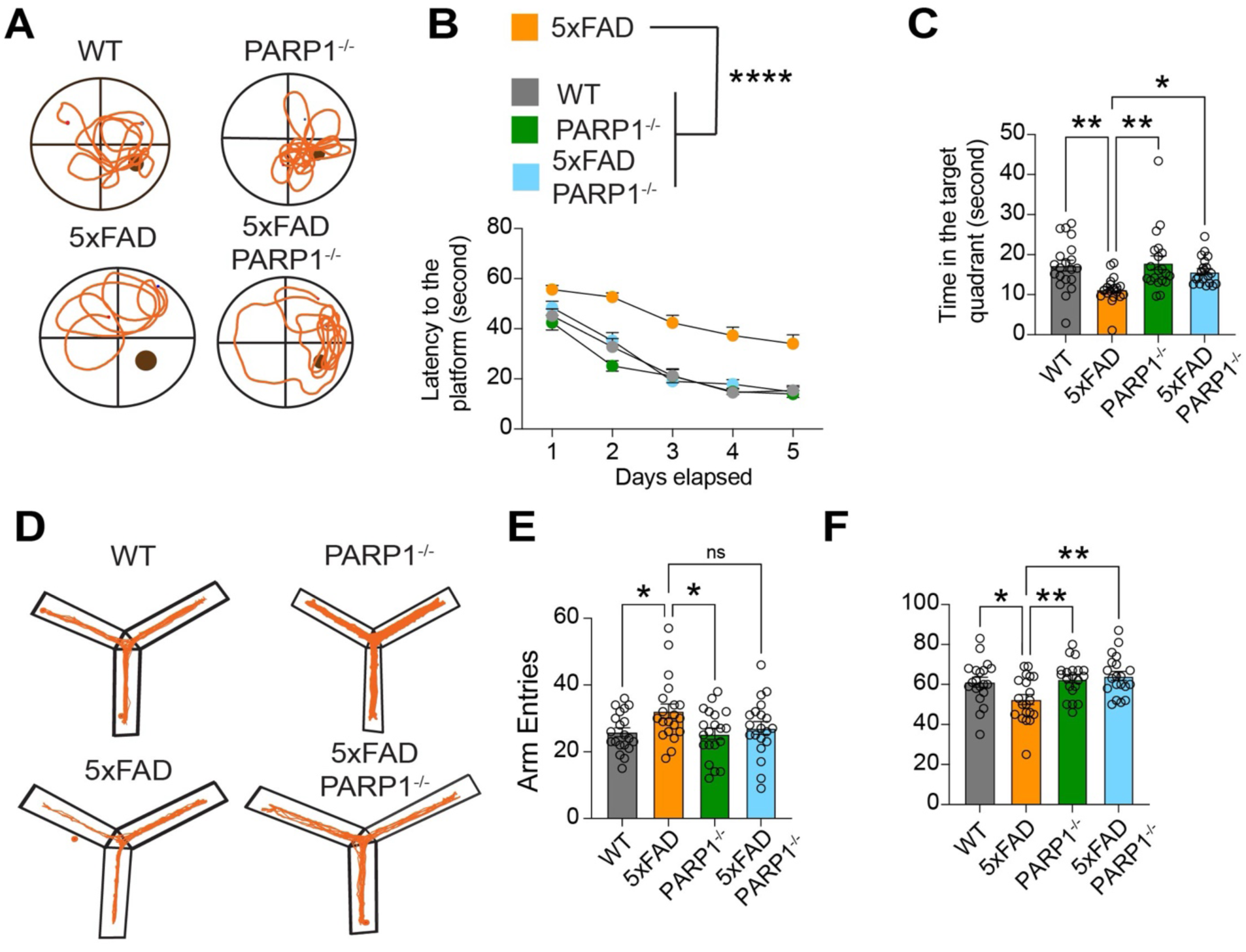
PARP1 deletion improves long term and spatial memory in 5XFAD mice. A, B and C) Morris Water Maze test data. A) Track plot of WT, PARP1^-/-^, 5XFAD and 5XFAD/PARP1⁻/⁻ mice on the day of probe trial. B) Latency to find the platform during training period. Bars are means ± SEM. Two-way ANOVA followed by Tukey’s post hoc test (n = 17-20). C) Probe trial data: time in the target quadrant. Bars are means ± SEM. Two-way ANOVA followed by Tukey’s post hoc test (n = 20). D, E and F) Y-Maze test. D) Trackplot of WT, PARP1^-/-^ , 5XFAD and 5XFAD/PARP1⁻/⁻ mice. E) Number of arm entries. F) Percentage alterations. Bars are means ± SEM. Two-way ANOVA followed by Tukey’s post hoc test (n = 20).

### The absence of PARP1 impacts amyloid precursor protein (APP) levels

Given that Aβ_1-40_ and Aβ_1-42_ levels were reduced, while APP mRNA levels remained unaltered upon PARP1 depletion in 5XFAD mice, APP protein expression and processing were subsequently assessed. Human full length APP ELISA revealed a decrease in APP levels in 5XFAD/PARP1^-/-^ mice (Fig. 7A). Additionally, there were similar statistical differences in full length APP levels and its processed byproduct, APP-CTF levels in 5XFAD/PARP1^-/-^ mice (Fig. 7*B-D*). Since, APP mRNA levels showed no statistical difference in 5XFAD and 5XFAD/PARP1^-/-^ (Fig. S2*D*), the observed differences in APP/APP-CTF levels were postulated to result from alterations in proteins involved in APP processing and in Aβ degradation.

**Figure 7.**
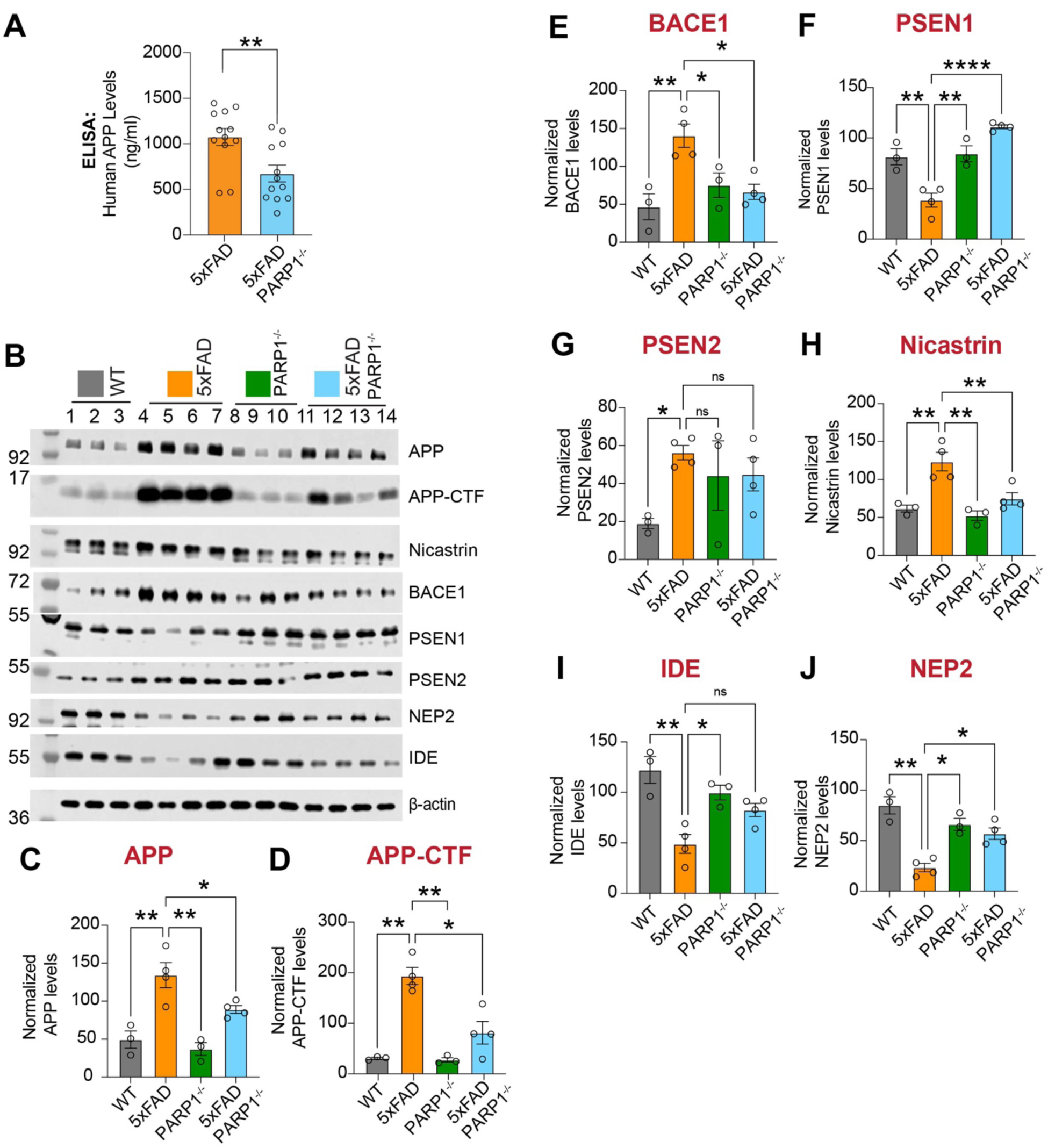
PARP1 deletion impact APP metabolism 5XFAD mice. A) ELISA quantification of APP levels in 5XFAD and 5XFAD/PARP1⁻/⁻ mice. Bars are means ± SEM. Unpaired Student’s t test (n = 12). B-J) Immunoblot and quantification of APP, APP-CTF and APP metabolism enzymes: Beta-secretase 1 (BACE1), Presenilin1 (PSEN1), Presenilin1(PSEN2), Nicastrin, Insulin degrading enzyme (IDE) and Neprilysin(NEP2). Bars are means ± SEM. Two-way ANOVA followed by Tukey’s post hoc test (n = 3).

5XFAD/PARP1^-/-^ mice showed a significant decrease in BACE1 protein levels (Fig. 7B, *E*) and BACE1 mRNA levels (Fig. S4*A*). Amongst the components of γ-secretase complex, PSEN1 protein levels were upregulated (Fig. 7*F*) and nicastrin protein levels were reduced (Fig. 7B, *H*), while PSEN2 protein levels remained unchanged (Fig. 7B, *G*). At the mRNA level PSEN1, PSEN2 and nicastrin showed no significant differences (Fig. S4*B-D)*. Enzymes involved in Aβ degradation, such as Neprilysin (NEP2) was upregulated in 5XFAD/PARPP1^-/-^ mice, while the insulin-degrading enzyme (IDE), remained unchanged (Fig. 7B, *I, J*). No statistical changes in mRNA levels were observed for NEP2 and IDE (Fig. S4*E, F)*. In summary, these results suggest that a reduced APP processing rate combined with enhanced Aβ degradation is a potential mechanism driving the decreased Aβ plaque load in 5XFAD/PARP1^-/-^ mice.

## Discussion

The major findings of this study demonstrate that PAR levels are significantly elevated in the CSF of patients with MCI and AD, with PAR levels negatively correlating with the Aβ_42/40_ ratio, a hallmark of amyloid pathology (24). Mechanistic investigations revealed that oligomeric Aβ_1-42_, but not Aβ_1-40_, activates PARP1 in primary cortical neurons, resulting in increased PAR synthesis, DNA damage, and neuronal death— effects that are prevented by both pharmacological inhibition and genetic ablation of PARP1. *In vivo*, genetic deletion of PARP1 in the 5XFAD mouse model of AD led to a reduction in amyloid plaque burden, preservation of synaptic density, prevention of neuronal loss, and significant attenuation of glial activation and neuroinflammation.

Importantly, PARP1 deficiency also rescued learning and memory deficits in these mice. At the molecular level, PARP1 ablation resulted in reduced BACE1 expression, altered γ-secretase complex composition, and upregulation of the Aβ-degrading enzyme NEP2, collectively shifting APP metabolism toward decreased Aβ production and enhanced clearance (25, 26). These integrated results identify PARP1 as an important modifier of amyloid-driven neurotoxicity, neuroinflammation, and cognitive decline in AD, and highlight its potential as a disease-modifying therapeutic target.

This study provides evidence that PAR levels are significantly elevated in the CSF of patients with MCI and AD, establishing a biomarker link between PAR metabolism and AD progression. The negative correlation between CSF PAR levels and the Aβ_42/40_ ratio in MCI and AD patients further supports the association of PAR with disease severity and amyloid pathology. Mechanistically, our *in vitro* data demonstrate that oligomeric Aβ_1-42_, but not Aβ_1-40_, activates PARP1 in primary cortical neurons, leading to increased PAR synthesis and DNA damage, as indicated by γ-H2AX elevation. Pharmacological inhibition or genetic ablation of PARP1 confers significant neuroprotection against Aβ_1-42_ toxicity, implicating parthanatos, in Aβ-induced neurodegeneration supporting prior findings. Consistent with these findings is the observation that PARP inhibition protects against Aβ peptide toxicity (27–30)

In vivo, 5XFAD transgenic mice lacking PARP1 exhibit a reduction in amyloid plaque burden, preservation of synaptic density, and prevention of neuronal loss, particularly in brain regions most vulnerable to amyloidosis. These neuropathological improvements are paralleled by attenuated glial activation and neuroinflammatory marker expression, suggesting that PARP1 contributes to both neuronal and glial pathology in AD (31). Behavioral assessments further reveal that PARP1 deficiency rescues spatial learning, memory, and anxiety-like deficits in 5XFAD mice, underscoring the functional relevance of these molecular and cellular changes.

At the mechanistic level, PARP1 ablation alters amyloid precursor protein (APP) metabolism by reducing BACE1 (β-secretase) expression and modifying γ-secretase complex composition, thereby decreasing Aβ production. In addition, PARP1 knockout enhances Aβ clearance through upregulation of NEP2, an Aβ-degrading enzyme, without affecting IDE. These findings suggest that PARP1 orchestrates both amyloidogenic processing and clearance pathways, and its loss shifts the balance toward reduced amyloid accumulation. The observed changes in BACE1 and γ-secretase subunits are consistent with PARP1’s known roles in regulating stress and inflammatory signaling, particularly via NF-κB, which can drive BACE1 transcription in response to Aβ-induced oxidative stress (28, 32).

PARP1 deficiency led to coordinated changes in the γ-secretase complex and Aβ clearance pathways that together may help explain the reduction in amyloid burden observed in 5XFAD/PARP1^⁻/⁻^ mice. Specifically, we observed an increase in PSEN1 protein, the primary catalytic subunit of γ-secretase, alongside a significant downregulation of nicastrin at both the protein and mRNA level, while PSEN2 protein remained unchanged despite reduced transcript levels. Since PSEN1 is essential for most Aβ generation (33), its upregulation may reflect a compensatory response to maintain γ-secretase activity; however, the concomitant loss of nicastrin—critical for substrate recognition—would be expected to impair complex assembly and function, ultimately resulting in reduced Aβ production despite increased PSEN1. This is further complemented by the observed decrease in BACE1, collectively shifting APP processing toward a less amyloidogenic route. In parallel, the absence of PARP1 enhanced Aβ clearance, as evidenced by increased an Aβ-degrading enzyme, NEP2 protein—a key Aβ-degrading enzyme—without changes in its mRNA, suggesting post-transcriptional regulation or stabilization (34). Given that NEP2 activity is known to mitigate amyloid pathology, its upregulation in PARP1-deficient animals would further accelerate Aβ catabolism. These integrated alterations—diminished amyloidogenic processing and enhanced degradation suggest how PARP1 ablation may orchestrate a multifaceted reduction in amyloid burden, extending prior evidence that chronic PARP1 activation exacerbates amyloid pathology and cognitive decline, while its inhibition confers protection in Alzheimer’s models.

Collectively, these results suggest that PARP1 may play an important role in mediating Aβ-driven neurotoxicity, amyloid pathology, neuroinflammation, and cognitive impairment in AD models. The integration of human biomarker data, mechanistic cellular studies, and in vivo genetic evidence supports further investigation of PARP1 as a potential target for disease-modifying therapeutic strategies in AD.

## ACKNOWLEDGEMENTS

This work was supported by grants from the NIH NS067525, AG085688, U19AG033655, the Thome Memorial Foundation, and the Alzheimer’s Association Zenith Award. T.M.D. is the Leonard and Madlyn Abramson Professor in Neurodegenerative Diseases. This manuscript is the result of funding in whole or in part by the National Institutes of Health (NIH). It is subject to the NIH Public Access Policy. Through acceptance of this federal funding, NIH has been given a right to make this manuscript publicly available in PubMed Central upon the Official Date of Publication, as defined by NIH.

## AUTHORS CONTRIBUTIONS

Conceptualization, A.J., T.-I.K., T.M.D., V.L.D.; Methodology, A.J., T.-I.K.; Validation, A.J., M.R.K., T.-I.K., M.R.K.; Formal Analysis, A.J., M.K., T.-I.K., M.R.K.; Investigation, A.J., M.K., T.-I.K., M.R.K., T.T., J.W., L.G., D.B., A.P., A.A., N.K., S.P., S.C, N.P.; Resources, A.M., M.A., L.M.B., J.B.L.; Writing-Original Draft, A.J., T.-I.K., T.M.D., V.L.D.; Visualization, A.J., M.K., T.-I.K.; Supervision, T.-I.K., T.M.D., V.L.D.; Funding Acquisition, T.M.D., V.L.D., A.M.

## DECLARATION OF INTERESTS

The authors declare no competing interests.

## Methods

### Animal

5XTg FAD (MMRRC Strain #034840-JAX) mice was obtained from the Jackson Laboratories (Bar Harbor, ME). The lab-maintained PARP1 ^-/-^ mice line was used to cross 5XTg FAD and PARP1^-/-^ together. After two generations, the desired littermates were aged and used for further experimentation. NIH Guide for the Care and Use of Experimental Animals and Johns Hopkins University Animal Care and Use Committee were followed for all housing, breeding, and subsequent procedures.

### Antibodies

Primary antibodies used for Western blotting and immunochemistry are listed in the Table below. These included antibodies against APP, APP-CTF, Nicastrin, BACE1, PSEN1, PSEN2, NEP2, IDE, β-Actin, IBA1, GFAP, Aβ (6E10 and 4G8), NeuN, RTN3, PSD95, PARP1, PAR, and γH2AX. All antibodies were obtained from commercial sources as detailed, except for the poly-ADP-ribose (PAR) antibody, which was prepared in-house.

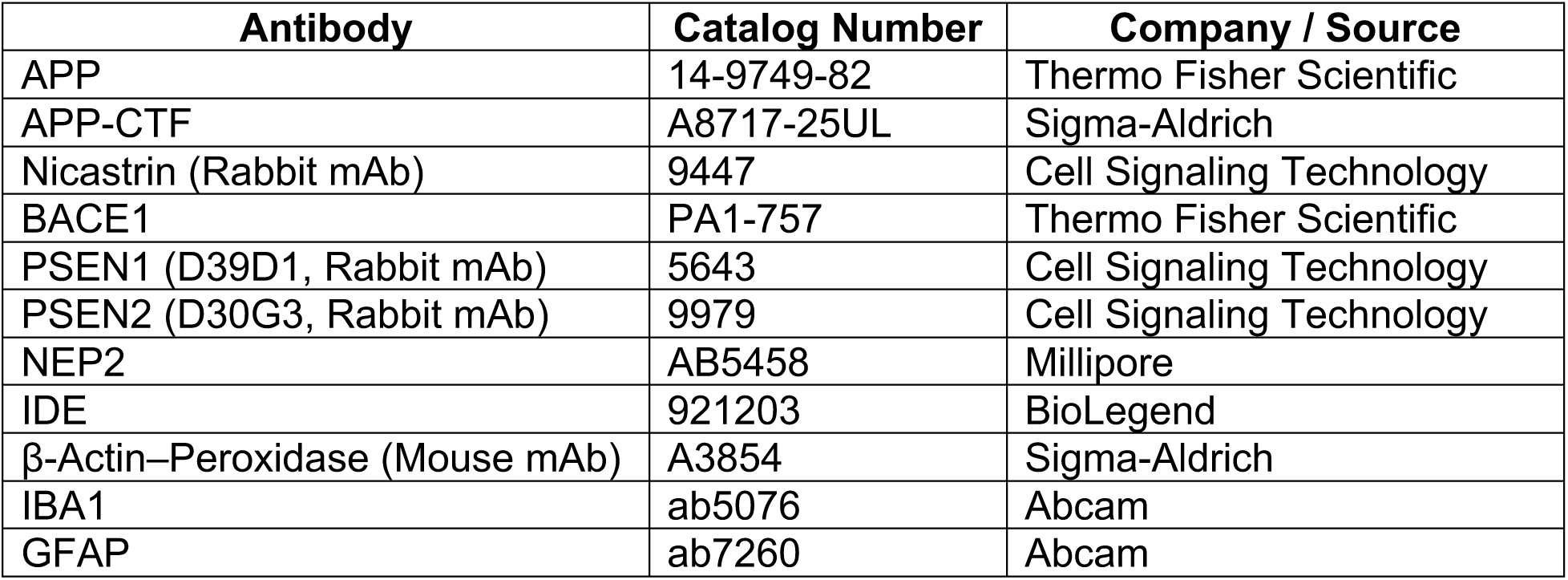
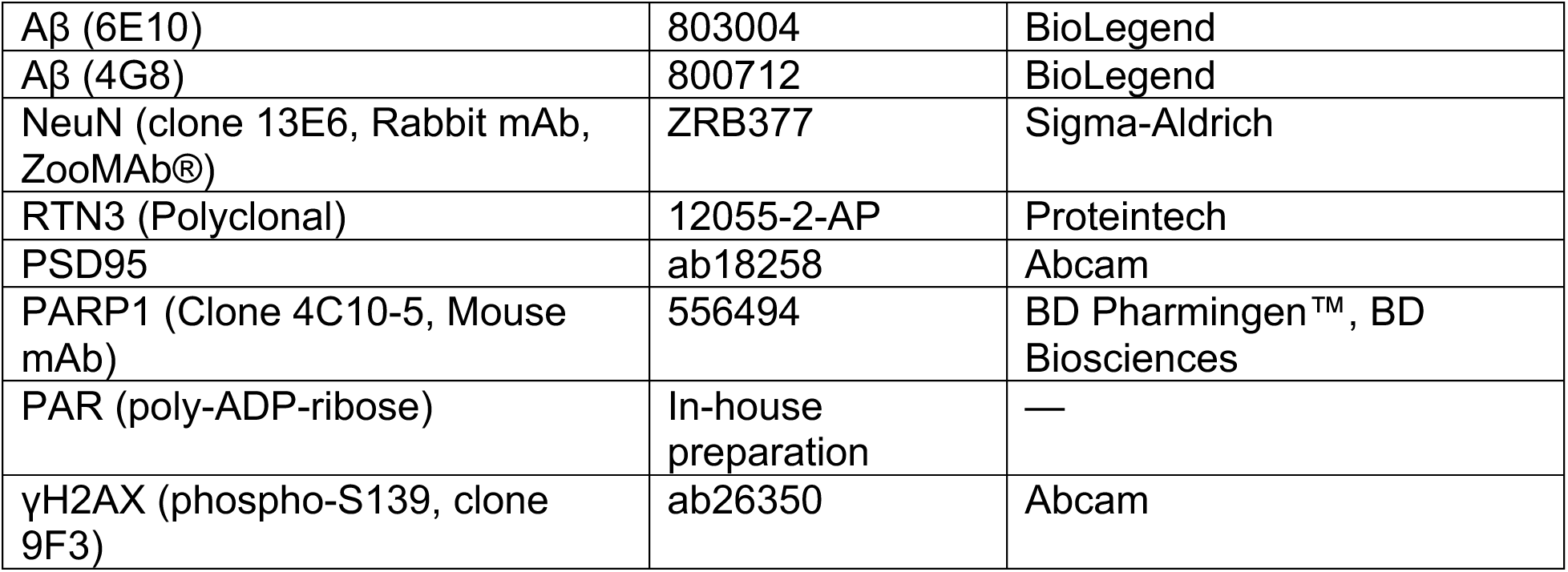

| Antibody | Catalog Number | Company / Source |
| --- | --- | --- |
| APP | 14-9749-82 | Thermo Fisher Scientific |
| APP-CTF | A8717-25UL | Sigma-Aldrich |
| Nicastrin (Rabbit mAb) | 9447 | Cell Signaling Technology |
| BACE1 | PA1-757 | Thermo Fisher Scientific |
| PSEN1 (D39D1, Rabbit mAb) | 5643 | Cell Signaling Technology |
| PSEN2 (D30G3, Rabbit mAb) | 9979 | Cell Signaling Technology |
| NEP2 | AB5458 | Millipore |
| IDE | 921203 | BioLegend |
| β-Actin–Peroxidase (Mouse mAb) | A3854 | Sigma-Aldrich |
| IBA1 | ab5076 | Abcam |
| GFAP | ab7260 | Abcam |
| Aβ (6E10) | 803004 | BioLegend |
| Aβ (4G8) | 800712 | BioLegend |
| NeuN (clone 13E6, Rabbit mAb, ZooMAb®) | ZRB377 | Sigma-Aldrich |
| RTN3 (Polyclonal) | 12055-2-AP | Proteintech |
| PSD95 | ab18258 | Abcam |
| PARP1 (Clone 4C10-5, Mouse mAb) | 556494 | BD Pharmingen™, BD Biosciences |
| PAR (poly-ADP-ribose) | In-house preparation | — |
| γH2AX (phospho-S139, clone 9F3) | ab26350 | Abcam |

### Human CSF samples and PAR ELISA

Participants in the Johns Hopkins University BIOCARD study (35) and at the Cleveland Clinic underwent annual evaluations, including detailed medical history, physical examination, and neuropsychological testing. CSF specimens were processed within one hour of collection: samples were centrifuged, divided into aliquots, and stored at – 80 °C at either the Cleveland Clinic Lou Ruvo Center for Brain Health Biobank or the Johns Hopkins University repository. For PAR quantification, two monoclonal anti-PAR antibodies (clones #19 and #25) were employed in an ELISA format. Ninety-six–well plates (NUNC, Cat. #46051) were coated overnight at 4 °C with clone #19 (5 µg/mL) as the capture antibody. Purified PAR standards (0–200 nM) and CSF samples from control or PD subjects were added and incubated for 1 hour at room temperature. Wells were washed five times with PBST (0.05% Tween-20 in PBS), followed by a 1-hour incubation with biotinylated clone #25 as the detection antibody. Signal development was achieved using HRP-conjugated streptavidin (Thermo Scientific), yielding a lower detection limit of ∼3 pM and saturation at 50 nM.

### Primary cortical neuron culture and treatment

Primary cortical neurons were isolated from embryonic day-16 WT or PARP1^-/-^ mouse embryos as previously described (5). Cells were seeded in Neurobasal medium supplemented with B-27, 0.5 mM L-glutamine, and 100 U/mL penicillin–streptomycin (Invitrogen, Carlsbad, CA). At DIV 7, cultures were pretreated for 1 hour with one of the following: ABT-888 (veliparib) at 1 µM, AG-014699 (rucaparib) at 1 µM, BMN 673 (talazoparib) at 1 µM. Thereafter, 1 µM oAβ_1-42_ or 1 µM oAβ_1-40_ were added, and cells were incubated for the indicated durations before analysis by cell-death assays or biochemical methods.

### Synthetic oligomeric oAβ_1-42_ preparation

Synthetic Aβ_1-42_ oligomers were generated from lyophilized monomers (rPeptide, Bogart, GA). Briefly, HFIP-treated Aβ_1-42_ was first dissolved in DMSO, then diluted into PBS to the desired concentration. The solution was incubated at 4 °C for 24 h to allow oligomer formation and subsequently stored at –80 °C. Prior to use, samples were centrifuged at 12,000 × g for 10 min, and the cleared supernatant was collected as the oligomeric Aβ (ADDLs). Oligomerization was confirmed by Western blot analysis (36).

### Cell death and viability assessment

Primary cortical neurons were exposed to 1 μM oAβ_1–42_ (oAβ1–42) for 48 hours. PARP inhibitors, including Talazoparib (BMN 673; LT-673, Catalog No. S7048), AG-14361 (Catalog No. S2178), and Veliparib (ABT-888; NSC 737664, Catalog No. S100), were added 30 minutes prior to oAβ_1–42_ treatment. Cell death was quantified by co-staining with 7 μM Hoechst 33342 and 2 μM propidium iodide (PI) (Invitrogen), followed by automated image acquisition and analysis on a Zeiss microscope using Axiovision 4.6 software (Carl Zeiss, Dublin, CA). Subsequently, Alamar Blue reagent (Invitrogen) was added, and cell viability was measured fluorometrically (λ_ex_ 570 nm, λ_em_ = 585 nm) as described previously (37).

### Amyloid-β ELISA

The ELISA for ascertaining different amyloid species-Aβ1-42 (Thermo Catalogue-KHB3441) and Aβ1-40 (Thermo catalogue-Cat #KHB3481) were performed as per the kit instructions.

### Immunohistochemistry and immunofluorescence

All mice were perfused with PBS and dissected to preserve half the hemisphere of the brain for immunostaining and other half for western blotting. The half saved for immunostaining was fixed overnight with 4% PFA followed by transfer to 30% sucrose for cryoprotection, where the brains remained until sectioned. Sample brains were sectioned at 50 μm thickness. Brains sections were then processed for immunostaining. By first incubating the sample in antigen retrieval buffer (Thermo catlog-00-4956-58).

This was followed by three PBS wash steps. The sections were then permeabilized using 0.3X triton X-100 contained in 10% goat serum. After this, the sections were blocked for an hour. Primary antibody incubation was performed overnight in a cold room. The next day, sections were PBS washed three times before incubating with secondary antibody for an hour at room temperature. After three washes, the samples were mounted using a mounting media containing DAPI. Similar steps were followed for immunofluorescence of primary neurons. For the Thioflavin S staining procedure, each brain section was treated with a 500 µM solution of Thioflavin S (ThS, Sigma-Aldrich, USA) in 50% ethanol for a duration of 7 minutes followed by ethanol washes and were subsequently mounted using mounting media. For Nissl staining, sections were counterstained with Nissl (0.09% thionin). The cell counting was performed using stereo investigator software. Immunofluorescence imaging was performed using confocal microscope-LSM880. Signal intensity and plaque counting was performed using ImageJ software (Kam et al. 2018).

### Real-Time Quantitative PCR

For the real-time quantitative PCR (RT-qPCR) procedure, total RNA was isolated utilizing TRIzol, following the manufacturer’s protocol. One microgram of the extracted RNA underwent reverse transcription with the High-Capacity cDNA Reverse Transcription Kit. Gene expression analysis was performed using SYBR Green-based RT-qPCR on an ABI ViiA 7 system (Applied Biosystems, Foster City, CA, USA), with results normalized to GAPDH levels. The following forward (F) and reverse (R) primers were employed (5′ to 3′, human sequences unless stated otherwise):

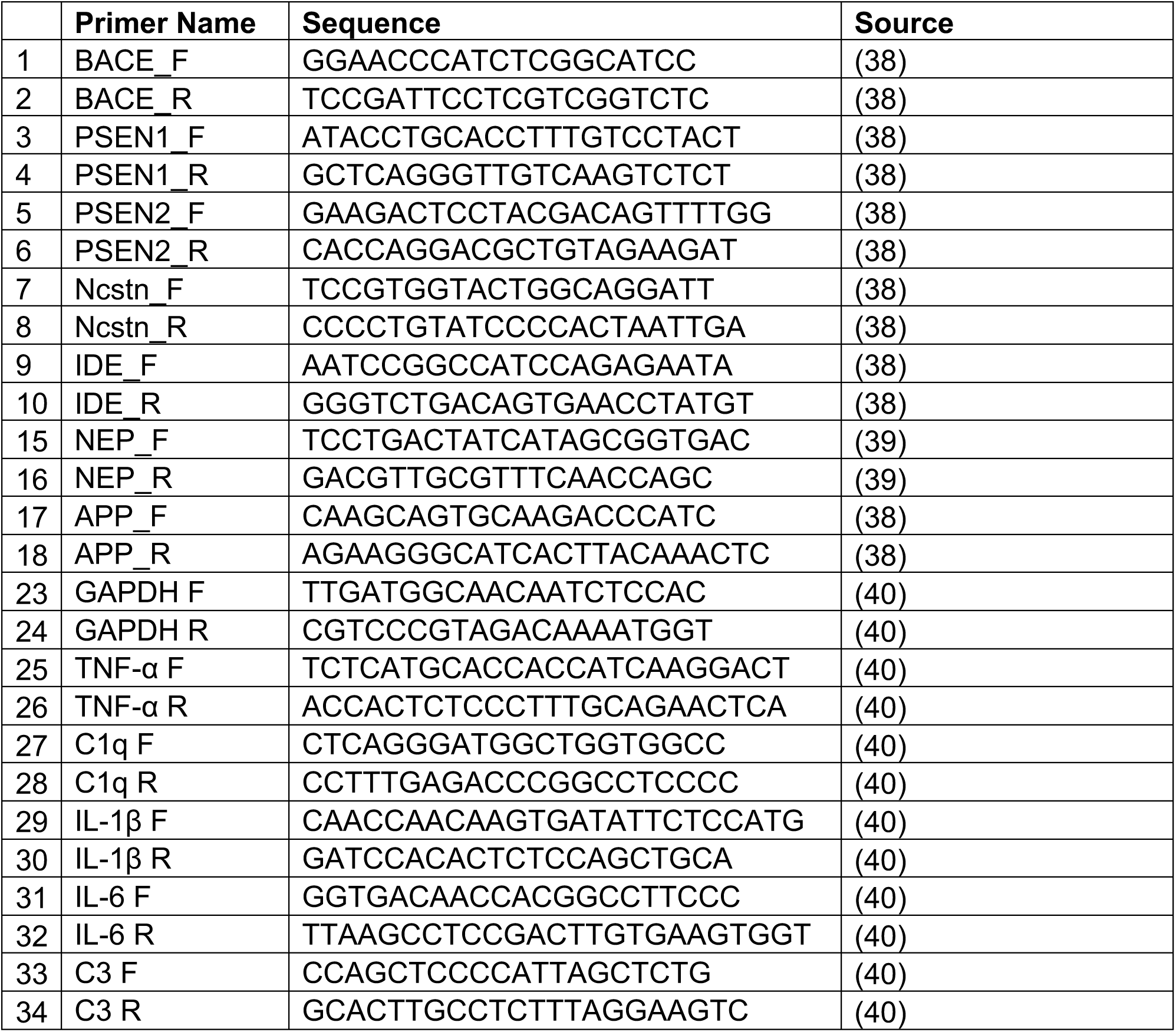

### Behavioral tests

#### Morris Water Maze

The Morris water maze was performed as published by (Park et al. 2021) with minor modifications. A circular pool (120 cm in diameter and 35 cm in height) was filled with water containing a nontoxic water-soluble white dye. The pool was split into four equal quadrants. A platform (8 cm in diameter) was randomly placed in one of the quadrants such that it does not appear visible (1 cm below the water surface), possibly in the center of that quadrant. Visual cues were attached around the quadrants to act as spatial references. A day before the trials mice were given a 60s swimming training in the absence of a platform. This is followed by a five-day training period wherein the mice swam three time (trials) a day, with an intertrial interval of 30 minutes, to develop memories to allocate the hidden platform, termed as escape latency. During this period, a mice was given 60s to find the platform, if successful within this time frame the test automatically ends, but if they are unsuccessful, they are made to stand on the platform for 10s. On the final day-probe trial, the platform was removed. The mice were again given 60s to swim. The time and distance spent in the target quadrant (previously containing the platform) was recorded. For the entirety of this experiment, ANY-maze software (ANY-maze system, Wood Dale, IL, USA) was used for recording.

#### Y-maze Spontaneous Alteration

The Y-maze spontaneous alteration was performed as published by (Park et al. 2021) with minor modifications. Mice were given five minutes to freely explore a Y-maze (40 × 8 × 15 cm). Using ANY-maze software the number of arm entries and percent alternations by the mice were recorded. An entry was considered legitimate only when all four limbs of the mice were within the arm.

#### Open field test

The Open field test was performed as described (Kim et al. 2019). Using Photobeam Activity System/PAS software (SD instruments), a rectangular box (40 cm x 40 cm x 40 cm) was digitally sub divided into 36 (6 x 6) identical sectors (6.6 cm x 6.6 cm), which was further subdivided into peripheral and central sectors. The mouse was placed inside this box in the dark and its movement was monitored via software for 30 minutes (5 minutes each six cycles). Between mouse change, the apparatus was thoroughly cleaned using Vinoba. The time spent in periphery versus the center was collected and graphed as a marker for anxiety.

#### Tissue Immunoblot Analysis

Frozen tissue from dissected brains were homogenized in Tris-buffered saline (TBS) homogenization buffer (20ul/mg). The sample was centrifuged at 100,000 × g for 1 h at 4° C using Optima TLX Ultracentrifuge (Beckman Coulter). The supernatant was transferred to a prechilled Eppendorf and pellet was resuspended in TBS buffer containing 1% Triton X-100 (20ul/mg). This time the sample was sonicated and then centrifuged at 100,000 × g for 1h at 4°C using Optima TLX Ultracentrifuge. The supernatant was saved as TBSX soluble. The remaining pellet was finally extracted using 70% formic acid. Following similar rounds of sonication and centrifugation, supernatant was saved as 70% formic acid soluble. This protocol is adapted from (Youmans et al. 2011).

#### Aggregate Aβ extraction

Mouse cortical tissue was lysed using an ice-cold buffer consisting of 10 mM Tris-HCl (pH 7.5), 150 mM NaCl, 5 mM MgCl₂, 0.5 mM DTT, 100 µg/mL cycloheximide, along with protease and RNase inhibitors, and 0.05% sodium deoxycholate. The homogenization was performed with a Dounce homogenizer. The resulting homogenates were incubated at 4 °C for 20 minutes and then cleared through centrifugation at 10,000 rpm for 10 minutes at 4 °C. The supernatants were collected, quantified using a BCA assay, and normalized for total protein content. A sucrose cushion was created by dissolving 2 g of sucrose in 4.7 mL of lysis buffer. For each sample, 900 µL of this cushion was layered beneath 600 µL of the normalized lysate, followed by centrifugation at 70,000 rpm for 2 hours at 4 °C. After ultracentrifugation, the supernatants were carefully removed, and the pellets were resuspended in 60 µL of lysis buffer. The enrichment of aggregates was verified through immunoblotting before proceeding to mass spectrometry.

#### Quantification and statistical analysis

Quantifications on immunoblots and immunofluorescent data was performed using ImageJ. For PSD95 density analysis, a Matlab script published in a previous paper was repurposed for analysis (41). Each color channel was first converted to binary signal.

The Aβ plaque image was used to outline the plaque using binary boundary detection. The code then dilated this boundary by 30 µm and applied the outlines onto the thresholded red channel to compute the area of the PS95 signal as a percentage of total area in each annulus.

All data are represented as mean ± s.e.m. At least 3 independent experiments were performed for *in vitr*o and immunofluorescence experiments. For behavioral experiments, the sample size (n) was about 20 mice. Statistical analysis was performed using GraphPad Prism 9 and CSF associated correlation analysis was performed using IBM SPSS. Differences between 2 means were calculated using an unpaired two-tailed student t test, and among multiple means using ANOVA followed by Tukey’s post hoc test. All p-values from the datasets were consolidated into a single table (Table S2).

## Supplemental Information

**Figure S1.**
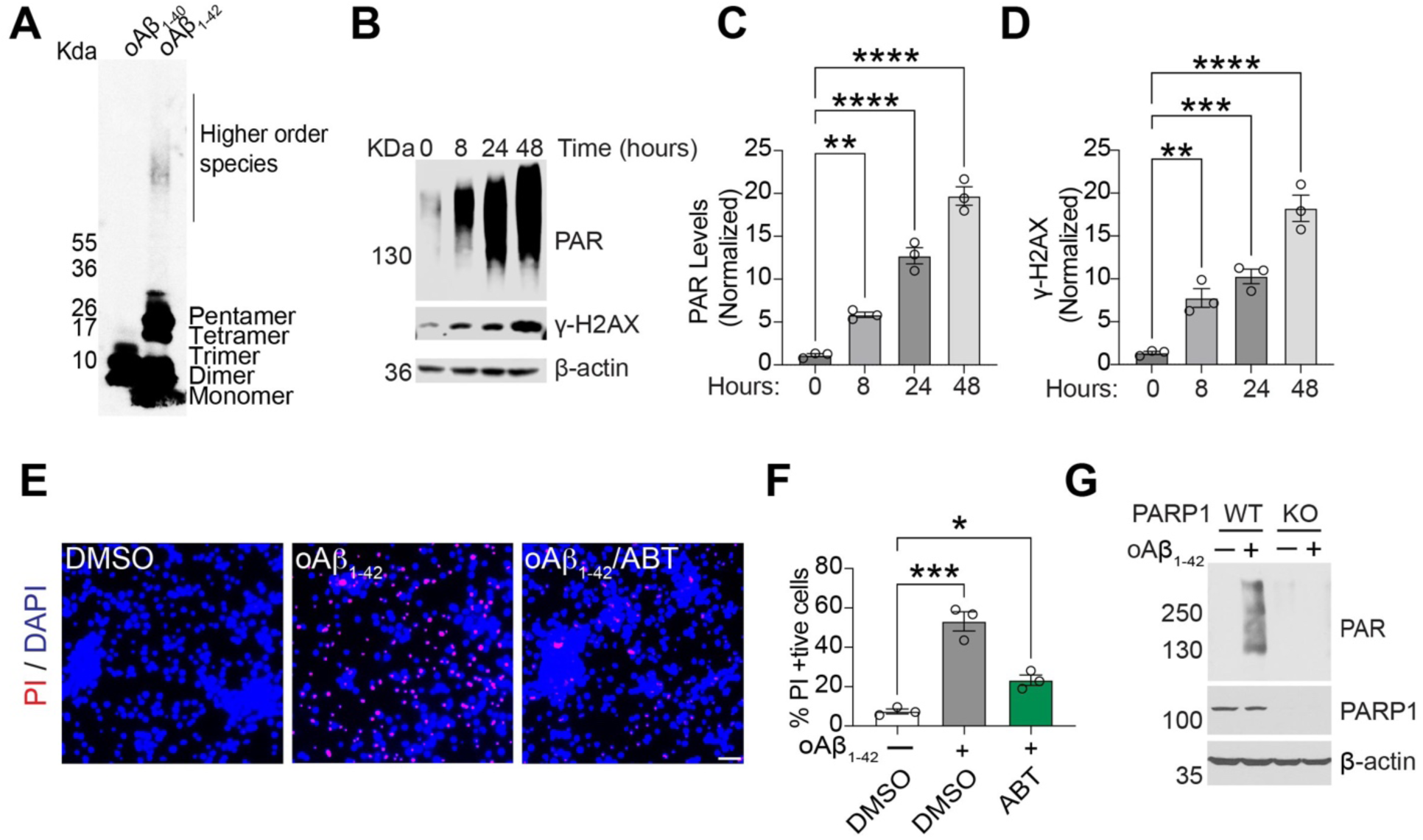
oAβ_1-42_ causes time dependent increase in PAR levels. A) Aβ_1-40_ and Aβ_1-42_ oligomers visualized using immunoblot. B-D) PAR and γ-H2AX levels in primary neurons treated with 1 µM oAβ_1-42_. (B) Representative immunoblot of PAR and γ-H2AX at the indicated time points. (C) Quantification of PAR levels over time. (D) Quantification of γ-H2AX levels over time. Bars represent mean ± SEM (n = 3). One-way ANOVA with Tukey’s post hoc test. E,F) 1 µM ABT888 protects against oAβ_1-42_ induced neurotoxicity. (E) Representative images of DAPI and propidium iodide (PI) staining from primary cortical neurons pre-treated with ABT-888 for 1 h, followed by further incubation with oAβ_1-42_ for 2 days. (F) Quantification of cell death. Bars represent mean ± SEM. Two-way ANOVA followed by Tukey’s post hoc test (n=3). G) Inhibition of PAR accumulation in primary cortical neurons of PARP1^-/-^ versus WT neuronal culture.

**Figure S2.**
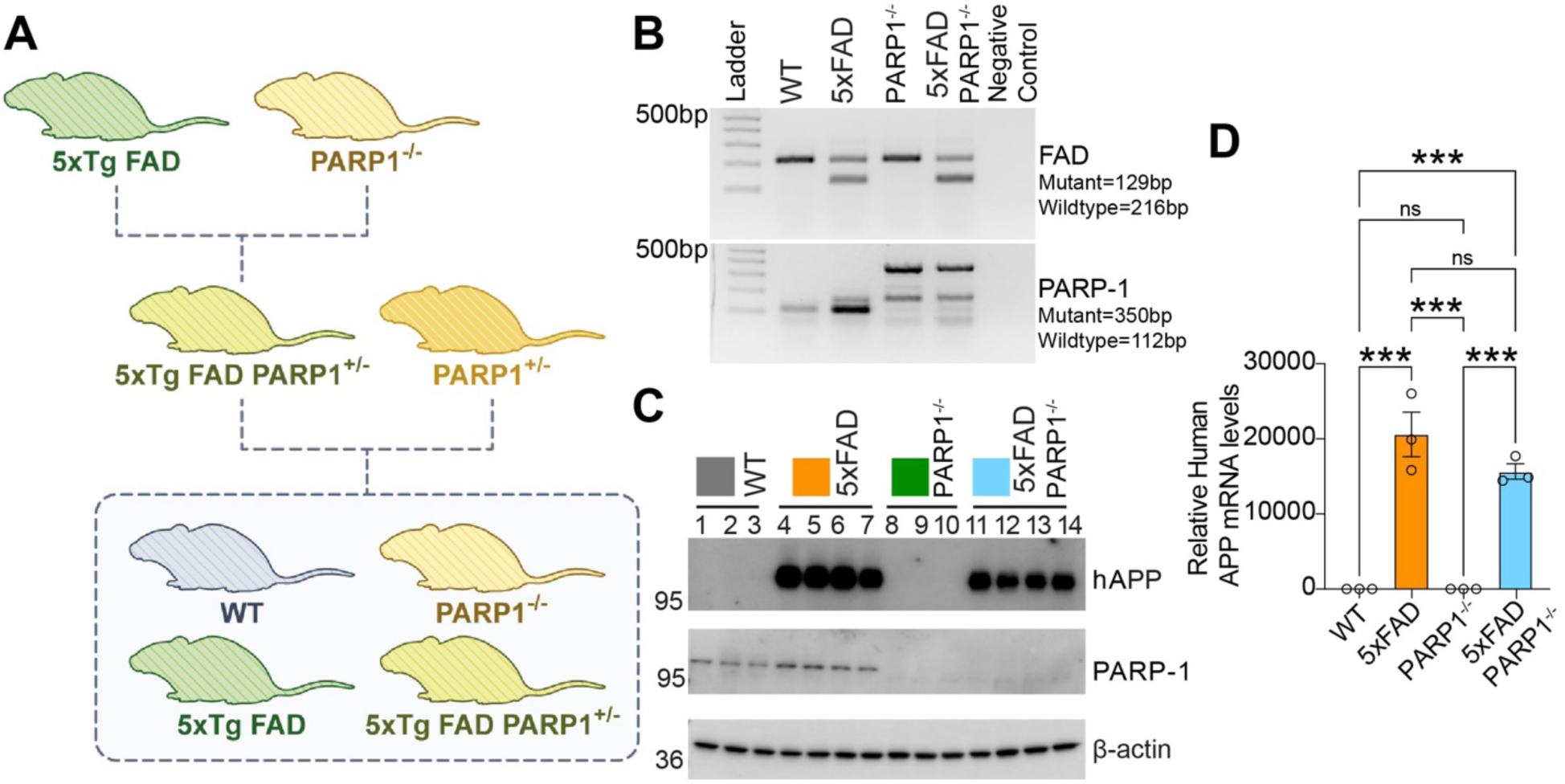
Generation of 5XFAD/PARP1^-/-^ mice. A-C) Breeding PARP1^-/-^ with 5XFAD mice. Data is shown along with (B) representative genotyping data verification and (C) immunoblot data. Genotyping was performed using primers sequence provided by Jackson labs. D) Human APP mRNA levels in 5XFAD and 5XFAD/PARP1^-/-^ mice. RT-PCR was performed to quantify human APP mRNA. Bars represent mean ± SEM. Two-way ANOVA followed by Tukey’s post hoc test (n=3).

**Figure S3.**
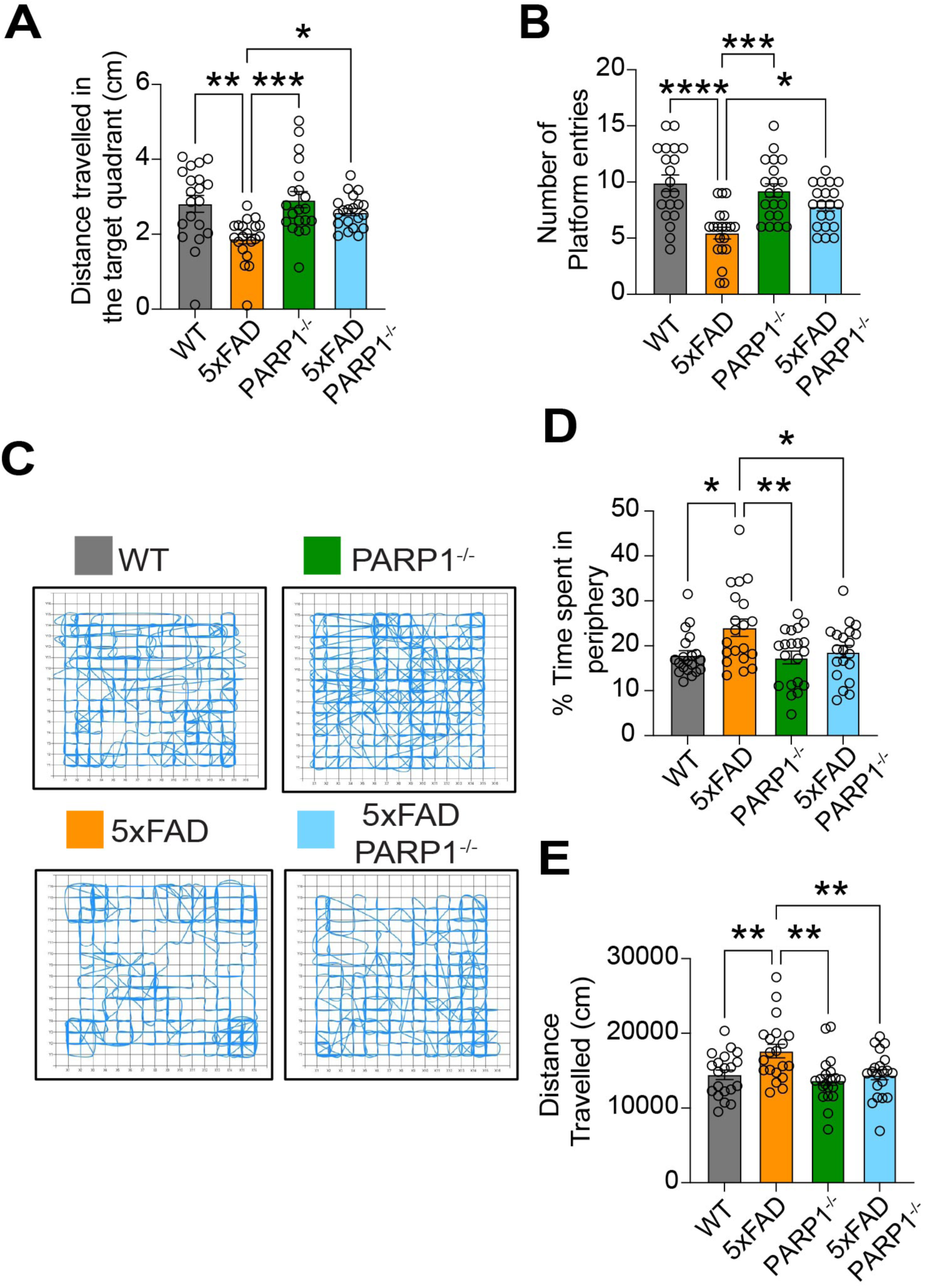
PARP1^-/-^ reduces anxiety like behavior in 5XFAD mice. A,B) Additional MWM Probe trial data: A) distance in the target quadrant and B) number of platform entries. Bars are means ± SEM. Two-way ANOVA followed by Tukey’s post hoc test (n =20). C-E) Open field test. C) Trackplot of WT, PARP1^-/-^, 5XFAD and 5XFAD/PARP1^-/-^ mice. D) Percentage of time spent in the periphery E) Distance travelled by the mice in open field test. Bars are means ± SEM. Two-way ANOVA followed by Tukey’s post hoc test (n = 20).

**Figure S4.**
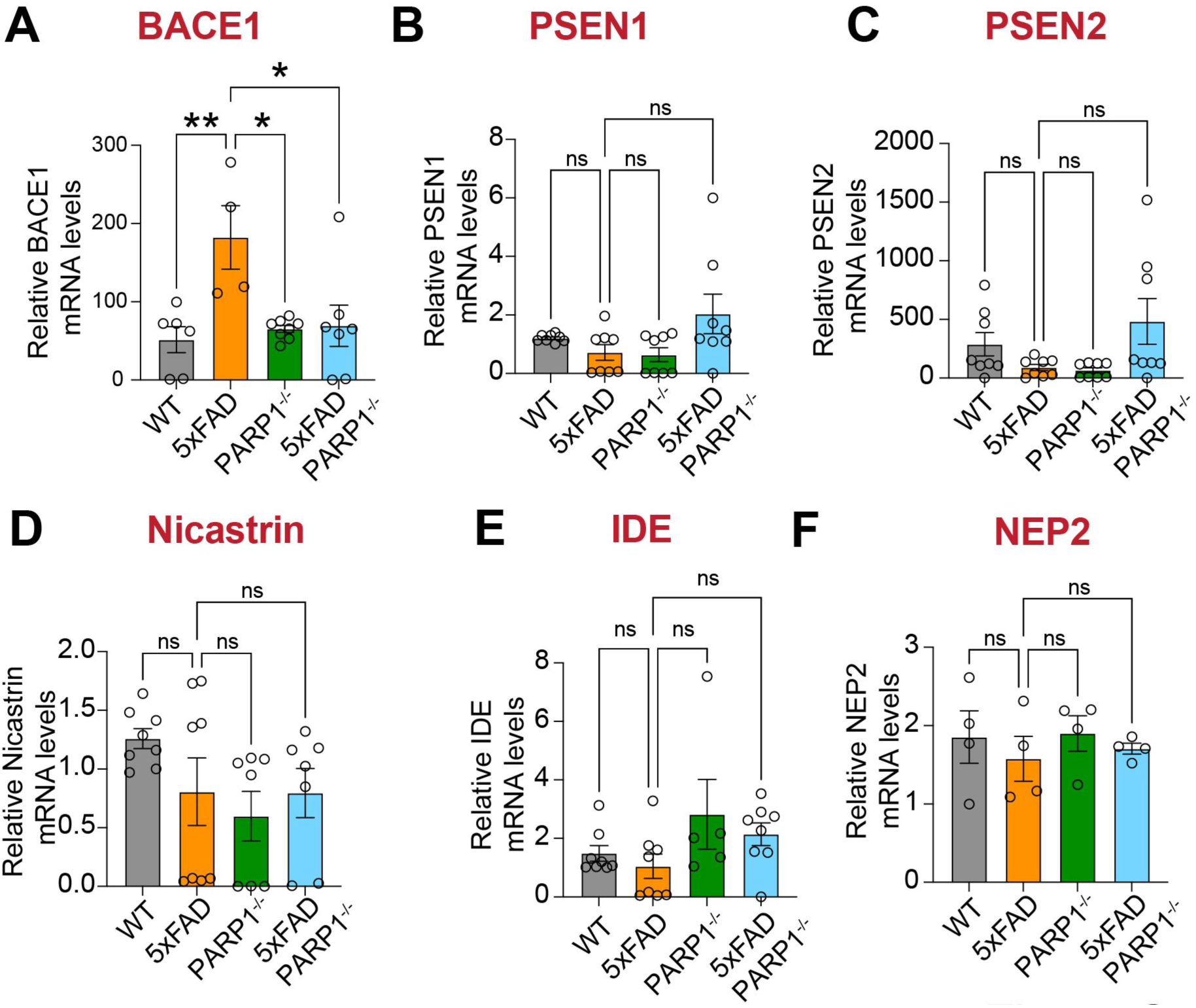
mRNA levels changes caused by PARP1 deletion in APP processing and degrading enzymes. A-F) mRNA levels of BACE1, PSEN1, PSEN2, Nicastrin, IDE and NEP2 as determined by RT-PCR. Bars represent mean ± SEM. Two-way ANOVA followed by Tukey’s post hoc test (n=5).

## Notes

### Competing Interest Statement

The authors have declared no competing interest.

